# Structural Basis of Voltage-Dependent Gating in BK Channels and Its Coupling to the Calcium Sensor

**DOI:** 10.1101/2023.12.29.573674

**Authors:** Gustavo F. Contreras, Rong Sheng, Ramon Latorre, Eduardo Perozo

## Abstract

The allosteric communication between pore domain, voltage sensors, and Ca^2+^ binding sites in the Ca^2+^-activated K^+^ channel (BK) shapes its multiple physiological roles as the preeminent signal integrator in excitable systems. BK displays shallow voltage sensitivity with very fast gating charge kinetics, yet little is known about the molecular underpinnings of this distinctive behavior. Here, we explore the mechanistic basis of coupling between voltage-sensing domains (VSDs) and calcium sensors in *Aplysia* BK by locking the VSDs in their resting (R196Q and R199Q) and activated (R202Q) states, with or without calcium. Cryo-EM structures of these mutants reveal unique tilts at the S4 C-terminal end, together with large side-chain rotameric excursions of the gating charges. Importantly, the VSD resting structure (R202Q) also revealed BK in its elusive fully closed state, highlighting the reciprocal relation between calcium and voltage sensors. These structures provide a plausible mechanism where voltage and Ca^2+^ binding converge physically and couple energetically to define the conformation of the pore domain and thus, BK’ full functional range.

## Introduction

Physiological processes rely on feedback loops to maintain homeostasis in fluctuating metabolic conditions. The large conductance calcium- and voltage-activated K^+^ channel (BK) is essential in in establishing negative feedback loops to control the firing patterns of neurons, transmitter release, insulin secretion, and smooth muscle contraction ^1^. The unique feature of the BK channel lies in its ability to integrate membrane potential and intracellular calcium levels while deploying a large K^+^ conductance that efficiently controls cell excitability. BK channels comprise four identical α subunits (Slo1) ^2,3^ with seven transmembrane segments (TMD) and a large C-terminal cytosolic domain (CTD). A voltage sensing domain (VSD, segments S0-S4) and a pore-gate domain (segments S5-S6) reside in the membrane, adjacent to two homologous regulators of the conductance of K^+^ domains (RCK) in each subunit (RCK1 and RCK2), which assemble as tetrameric gating rings (4x CTD) and serve as the cytoplasmic Ca^2+^ sensor ^4^.

BK structure is highly correlated to its function in response to chemical stimuli ^5,6,10^. In BK, the closed to open transition (C→O) can be cooperatively facilitated by both intracellular Ca^2+^ and membrane potential ^11–13^. In addition, BK can be independently opened by Ca^2+^ binding^5–9^ and depolarizing voltage^12,14–17^ leading to near-maximal activation. BK channels display low probability openings at negative potentials without voltage or Ca^2+^ activation, resulting from independent transitions in the pore^12,15^. This gating behavior has been well described by allosteric models, such as the Horrigan-Aldrich (HA) model^12,15^, which describes the energetic interactions among sensors and gates as fluctuating between highly cooperative and independently coupled subunits. This coupling leads to simultaneous perturbations in the dynamics and equilibrium when individual sensors are active. Previous structural work probing Ca^2+^ activation demonstrated that the expansion of the gating ring in response to Ca^2+^ binding leads to conformational changes across all domains of the channel ^5–9^. Thus, BK voltage activation influences Ca^2+^ binding to the RCK1 domain^18^ triggering conformational rearrangements in the Ca^2+^ sensor^19,20^. Reciprocally, Ca^2+^ activation modulates the movements of VSD^21^ and shifts the equilibrium constant that define its resting-active transition^22^. However, the nature of BK voltage sensing, the strength of the VSD-pore interaction and its relevance to activation remains under debate ^15,22–24^.

Given BK’s dual activation regime, it should be possible to evaluate the major close-open state transitions by simply varying Ca^2+^ at a constant voltage. Indeed, a variety of BK cryo-EM structures have been obtained from channels in a variety of species^5,6,10,25–27^, under apo and Ca^2+^-bound conditions. Structures of Ca^2+^-bound BK channels at 0 mV satisfy the expectations of a fully open state, but analyses of Ca^2+^-free (closed) structures have been far less clear. This uncertainty stems from the fact that in the nominal absence of Ca^2+^, the intracellular gate remains sufficiently wide to allow a hydrated K^+^ ion or even large quaternary ammonium ions to pass through or access the inner water-filled cavity^28^, casting doubt on whether these structures truly represent closed conformations^29^. The HA allosteric model predicts that the channel may adopt intermediate closed states at 0 mV in the absence of Ca^2+^, where at least one voltage sensor is active^12,15^. Furthermore, large hydrophobic ions can block the channel in putatively closed states, yet fully closed BK, with all VSDs at rest, is not affected by these blockers^30^. This observation also raises questions regarding the positioning of the VSD charged residues in the apo structure, which point to only minor movements in S4, and no obvious rearrangements between the gating charges and their countercharge partners. To add to this uncertainty, the actual number of BK gating charges is still under scrutiny. To account for BK’s uniquely shallow voltage dependence and fast gating kinetics, Ma et al.^31^ proposed a decentralized voltage sensor where the gating charges are spread among S2, S3, and S4; while Carrasquel-Ursulaez et al.^32^, used gating current measurements to argue that only two arginines in S4 (R210 and R213) contributed to the gating charge. However, the nature of BK voltage sensing, the strength of the VSD-pore interaction and its relevance to activation remains under debate ^15,22–24^.

Here, by means of electrostatic engineering of the gating charges on the S4 helix, we investigated the effect of voltage on the BK’s channel conformations. Using single-particle cryo-EM, molecular dynamics simulations, and electrophysiological approaches to reveal the high-resolution structures of Aplysia BK underpinning its cooperative gating. We show that the nature and kinetics of VSD function in BK can be fully explained by a mechanism with limiting S4 helical excursions but large rotameric reorientations in the gating arginines. At the same time, the convergent coupling between VSD and Ca^2+^ sensing domain can be derived from the determination of a fully closed BK structure. These structures, in addition to those obtained with and without bound Ca^2+^, serve as a foundational framework for understanding BK allosteric interactions, and its role as signal integrator in excitable cells

## Results

### Neutralization of S4 charges polarizes the voltage sensor structure

We systematically neutralized the charged residues of the S4 (R196, R199, and R202) of the *Aplysia* BK (aBK) channel to evaluate their structural role in the channel activation (**Figure 1A**). These mutations were introduced in the construct previously used for structural determination of the *wild-type* (WT) structure of *aBK* ^5,6^. We recorded robust K^+^ currents from virus-infected sf9 cells for the WT and mutant aBK channels (**Figure 1B**). aBK VSD mutants produce significant shifts in the conductance-voltage relationship (G-V) similar to those found in mouse and human BK ^31–33^. The half-activation voltages (**V_0_**) of the mutants R196Q (−80 mV) largely leftward shifted, whereas R202Q shifted rightward relative to the WT(160 mV) (**Figure 1B**). These V_0_ shifts support the notion of a conserved voltage activation mechanism among different BK channel species.

**Figure 1.-.**
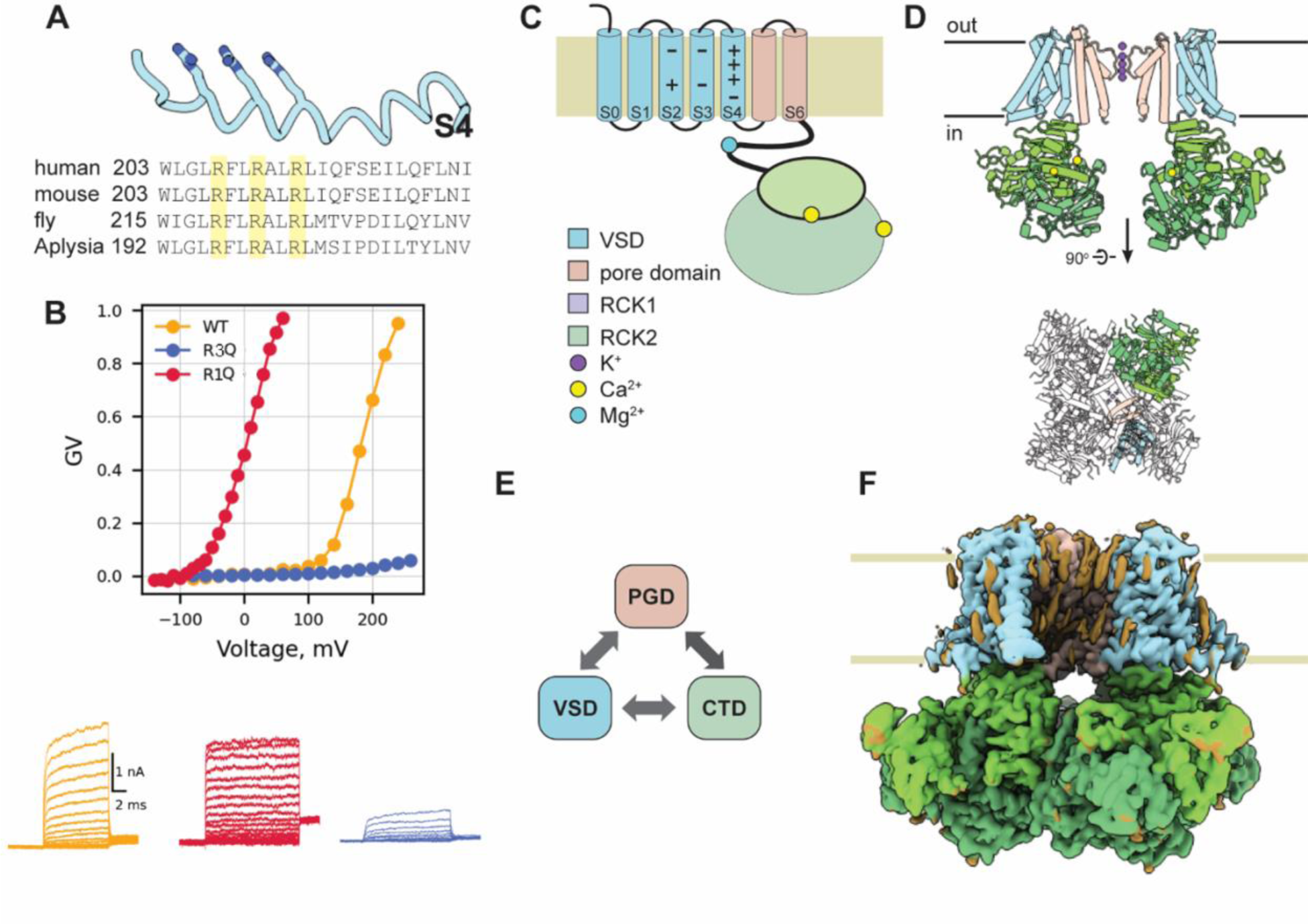
Function and architecture of *aplysia* BK. (**A)** Sequence alignment of BK S4 across different species. We focused on the mammalian arginine residues that modify the state of the VSD at 0 Ca^2+^. (**B)** *G*-*V* relations and representative traces aBK, aBKR196Q, and aBKR202Q. K+ current were recorded in sf9 infected cells under voltage clamp in whole-cell configuration; the internal solution contains 10 mM EGTA. (**C)** Schematic of aBK secondary structure features. The features are colored like (D-F). (**D**) Side and top views of the aBK structure are colored according to the different domains. Side view with only two subunits shown for clarity. (**E**) Schematic representation of interaction between different domains. (**F**) cryo-EM maps of aBK WT Ca^2+^-bound (WT-Ca).

We determined the structure of aBK mutant channels in detergent micelles using single-particle cryo-electron microscopy (cryo-EM). Specifically, we focused on three voltage-sensing arginines that are conserved in the BK S4, which we label as R1, R2, and R3. These residues correspond to positions 196, 199, and 202 in Aplysia, and to residues R207, R210, and R213 in human BK channel. For each of the mutants, two conditions were evaluated: the Ca^2+^-bound and apo conditions. The Ca^2+^-bound mutants were purified in solutions containing 10 mM Ca^2+^ and 10 mM Mg^2+^, while the apo condition purification was carried out in the presence of 2 mM EDTA. With the exception of R3Q-apo, all data sets produced one density with C4 symmetry. Although refinement with symmetry C1 and C2 were tested in the refinement, they resulted in models with lower resolution. The R3Q-apo data set resulted in two classes with C4 symmetry. The resolutions attained for each mutant were as follows: for R1Q, 2.7 Å in apo and 2.9 Å in Ca^2+^-bound conditions (**Figures S2A-B**); for R2Q, 3.4 Å in apo and 2.9 Å in Ca^2+^-bound conditions (**Figures S2C-D**); and for R3Q, 3.6 Å in apo and 2.6 Å in Ca^2+^-bound conditions (**Figures S2A-B**, Table S1). All structures display the overall topology characteristic of the WT BK channel. This includes a non-domain-swapped arrangement between the voltage-sensing domain (VSD) and the pore domain, alongside a domain-swapped cytosolic C-terminal regulatory domain (CTD) (**Figure 1C-F**). Comparative analysis of the refined atomic models (**Figure S3**) revealed that the positions of α-carbons from S4 arginine do not vary between Ca^2+^-bound and apo structures for all mutants (**Figure 2A**). Consistent with earlier data ^5,10^, we found that mutations in S4 stabilize the VSD structure solved in detergent micelles at 0 mV, independently of Ca^2+^. Nevertheless, we found variances in the structures of aBK mutants that shed light on the molecular mechanisms underlying channel gating.

**Figure 2.-.**
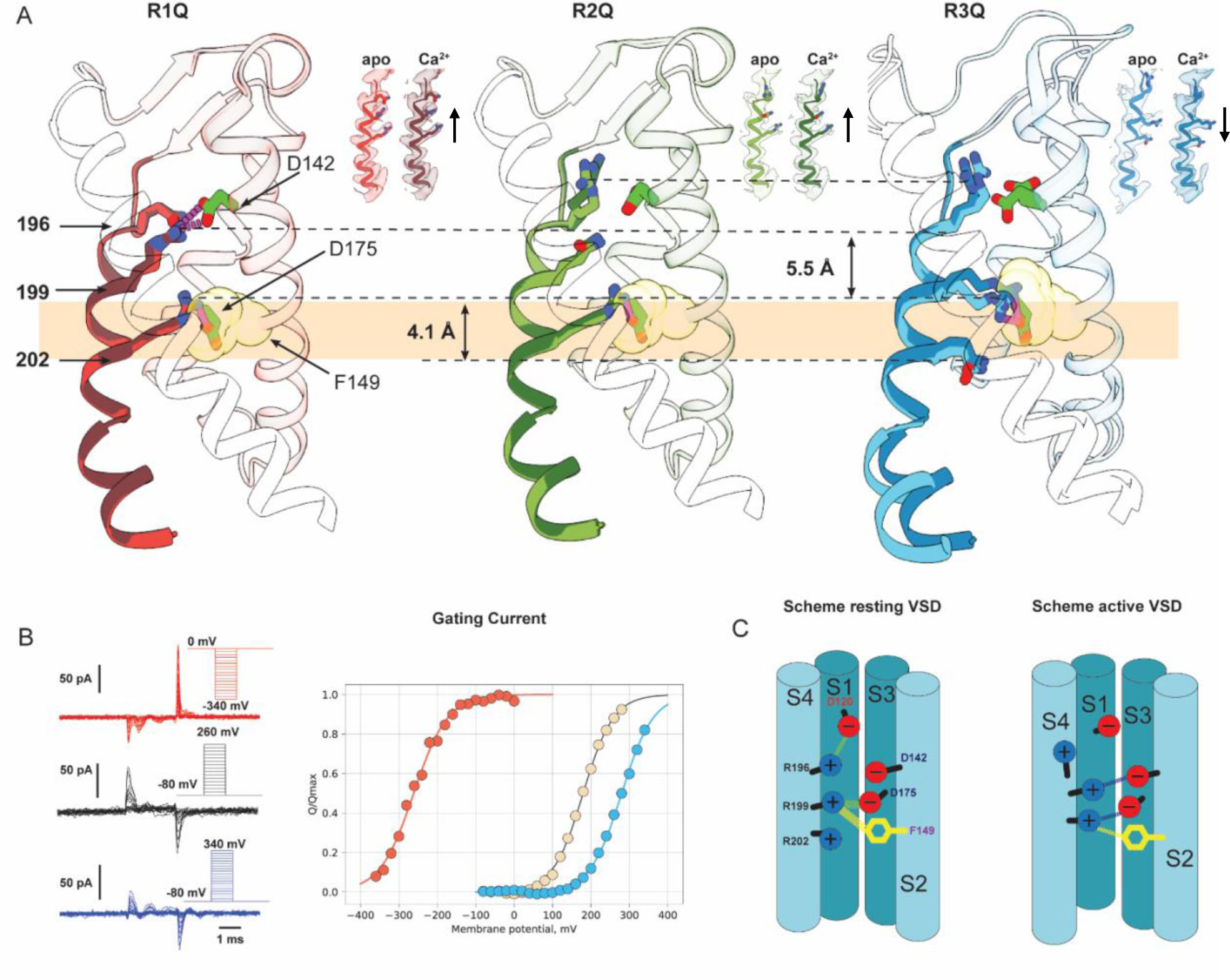
S4 charge neutralization stabilizes distinct VSD conformations independent of Calcium. **(A)** The VSD S4 of aBK mutants R1Q, R2Q and R3Q are shown as a cartoon, superimposed for condition Ca^2+^-bound and apo. For clarity, only S4 is shown as a cartoon, and TMD S1, S2, and S3 are shown in transparency. Stick side chains are shown for the charged residues: 196,199, 202. The structures were aligned based on residues of their selectivity filter 275-282 (S5-S6 loop, the least mobile part of *all*). **(B)** Representative traces of gating current and steady state Q-V curve for mutants R1Q (red), R3Q (light blue), and WT (beige). Q-V curves were obtained by 300 uS integration of the Qoff response. Half-activation voltage was obtained by adjusting data to a Boltzman distribution and the obtained values were WT 180 mV (n=4), R1Q (−230 mV, n=5) and R3Q 280 mV (n=6). The apo condition was achieved with 5 mM EGTA. **(C)** Schematic representation showing how the resting and active VSD conformational states.

### Structural Properties of the Activated Voltage Sensor

The activated state of the VSD was stabilized by neutralizing the first argine in the S4 segment of BK. We found that neutralizing R1 decreases the activation energy of VSD, as evidenced from ∼400 mV leftward shifts in the Q-V toward negative potentials (**Fig 2B**). This mirrors previous hBK studies, where neutralization of equivalent residues in hBK and mBK resulted in channels with facilitated activation but without changes in voltage sensitivity or coupling between VSD and the pore ^31,32^. When compared with the WT Ca^2+^-bound structure, R196Q shows essentially the same conformation at the TM region (segments S1 to S6, Cα RMSD < 1 Å)(**Fig S5**). In these R1Q structures, R2 forms hydrogen bonds with D142 (S2) at approximately 2.7 Å, while R3 interacts with F149 (3.2 Å) and D175 (3.2 Å) (**Figure 2A, Fig S4A**). These interactions are consistent not only in R196Q (both apo- and in Ca^2+^-bound) but also in the WT Ca^2+^-bound state (**Fig S4**). The introduction of glutamine at position 196 leads to a more acidic environment at the extracellular surface of the VSD, created by residues D120 and D142 (**figure S5**). This electrostatically negative environment draws in residues R2-R3, thereby stabilizing the active state of the VSD in the absence of Ca^2+^. Meanwhile, R1 cancel out the acidic environment the WT channel, allowing the VSD to adopt a resting conformation in the apo condition (**Figure 2E**). Thus, the structure confirms that R1 does not function as a gating charge but rather, it appears to modulate VSD transitions. This strongly suggests that the VSD in R1Q is equivalent to the active VSD in the WT channel.

The R1Q structures exhibit highly ordered TM domains, including the S0 helix. This helix was poorly resolved in previous detergent-based structures^5,6,10^, but in recent cryo-EM structures of hBK solved within native membranes, the S0 helix has been successfully delineated, and the higher resolution achieved may be explained by the stabilization effect of the membrane environment^25^. However, the present R1Q apo structure displays further detail in this region (**Fig S6),** allowing us to resolve five additional residues at the N-terminal end of S0. This higher structural order may be attributed to the non-canonical interaction between S4 and S0. Which involves aromatic residues in the extracellular end of S0^34^, that appear to be coupling this segment during the voltage activation^35^ and indicating that S0 helix is part of the voltage sensor.

As anticipated, voltage sensor activation opens the channel without calcium binding as shown in the R1Q apo structure **(Figure 3A**). Not only is the pore open but also the Ca^+2^ sensor of R1Q appears expanded in the apo condition despite the lack of strong density for Ca^2+^ at its binding site in the R1Q apo (**Fig S7**). In the WT channel, upon Ca^+2^ binding the CTD undergoes an expansion when compared to the WT apo condition (Cα RMSD ∼4 Å). We found minor differences between apo and Ca^2+^-bound conditions in the R1Q structures (Cα RMSD ∼1.6 Å), suggesting that the active VSD already drives a conformational change in CTD structure. This finding aligns with previous functional studies^19,21,22^, suggesting that a robust coupling between the VSD and CTD underlies the gating of the channel.

**Figure 3.-.**
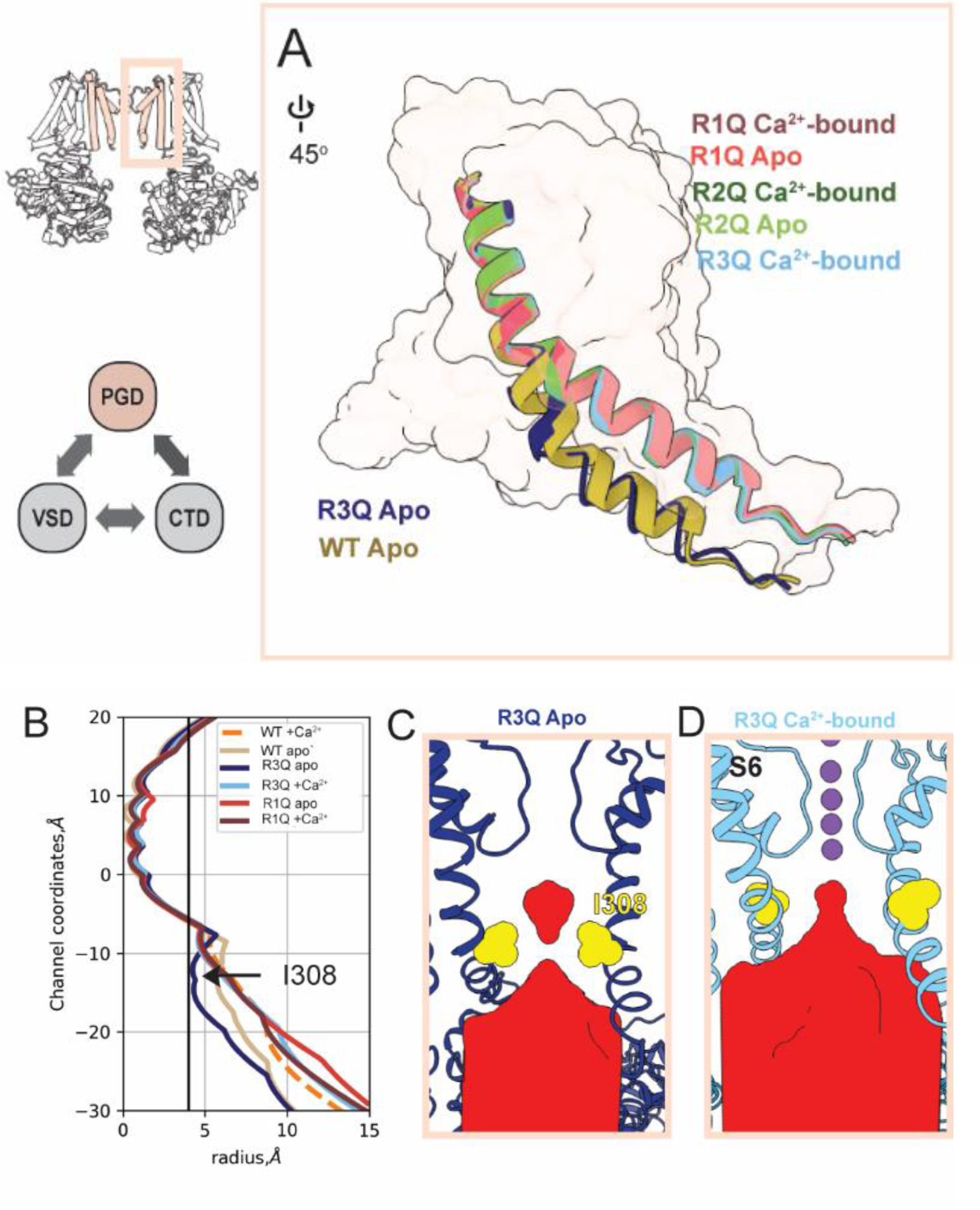
aBK deep close state revealed by R202Q-apo. (A) Superimposed stereo view of the pore lining segment S6 for WT-apo and mutants R1Q, R2Q and R3Q in both apo and Ca^2+^-bound conditions. For clarity, only S6 is shown as a cartoon representation, and the TMD is shown as a surface. (B) Pore-size profiles of the permeation pathway calculated by HOLE () (C) Side view of the pore, with only two subunits shown for clarity. The effective opening radius is represented by HOLLOW

R2Q also produced active-like VSD structures, regardless of Ca^2+^ binding (**Figure 2A**). However, R2Q VSD lacks coupling between S4 and S2 at residues R2-D124 as seen in the active VSD structures R2Q and WT (**Fig 2A**). This coupling has been tracked using fluorescent probes attached to the extracellular end of S4 and S2^36^. Neutralization of either R2 or D124 resulted not only in the loss of a voltage dependent S4-S2 coupling, but also it reflects an increase in the activation energy to open the channel^36^. This suggests that the interaction between S2 and S4 plays a pivotal role in both voltage sensitivity and the VSD-pore coupling. Structurally, R199 undergoes conformational changes during VSD activation, that are better highlighted by the in the resting VSD structure.

### Structural Properties of the Resting Voltage Sensor

The resting state of the VSD is stabilized by the neutralization of arginine at position 202, as predicted by a rightward shift of about 100 mV in the Q-V (**Figure 2B**). The charged residues of S4 (R1 and R2) maintain consistent positions in the R3Q-apo and R3Q Ca^2+^-bound structures. R1 (S4) forms salt bridges with D120 (S1) with a bond length of 2.7 Å, and R2 establishes hydrogen bonds with D175 (S3) and exhibits weak interactions with F149 (S2) (**Fig. 2A**). These two interactions contribute differently to channel gating. The interaction at the extracellular side of the VSD (R3-D120) has a modulatory role. The zone comprising the countercharge D175 (S3) and the aromatic ring F149 (S2) appears role in coordinating charge transition during VSD activation, it coordinates R2 in the resting and R3 in the active VSD structure, similar to the hydrophobic plug in Kv channels (**Figure 2A**). Neutralization of the primary countercharge in the resting VSD, hD186N (D175), reduces the coupling between VSD activation and channel opening, as described in previous^31,32^. The resting VSD structure (R3Q-apo) show that R2 undergoes conformational changes during VSD activation. In the resting VSD structure (R3Q-apo), R2 faces the intracellular solution, whereas in the active VSD state (R1Q-apo), it faces the extracellular solution (**Figure 2A**). In both cases, R2 forms obligatory interactions for channel function. This finding agrees with the functional characterization of residue R2, and confirm that R2 is the main gating particles of the BK VSD^32^.

Stabilizing the VSD in its resting (down) conformation leads to a fully closed pore in the apo condition, producing two structural classes with minimal changes in their TM region (**Fig S8**). Two classes were generated for R3Q-apo using 3D classification in cryosparcs, with overall resolutions ranging from 3.6 to 3.7 Å (**Figure S9**). These classes are related to each other by a small rotation of the transmembrane domain of up to 8° about the central four-fold axis of the channel, similar to the variations found in WT apo structures at different EDTA concentrations^5^. Both structures display minimal differences in the pore domain (Cα RMSD > 1 Å) and more notable differences in the VSD (Cα RMSD ∼ 1.5 Å) and the CTD (Cα RMSD ∼ 2.2 Å). In the R1Q-apo dataset, a greater degree of heterogeneity was observed, which may be attributed to structural disparities between *aplysia* and *human* BK, particularly in a cluster of positively charged residues known as the RKK ring. These residues encircle the entrance of the pore and are thought to stabilize the closed conformation due to their interaction with negative charges at S6^37^. In aBK, the second lysine in the ^329^RKK^331^ ring is neutralized (^318^RQK^320^). When this lysine is substituted with alanine in hBK, it results in a 100-fold increase in spontaneous activity at 0 mV, suggesting that aBK is inherently more disordered in the closed state.

### The resting VSD structure reveals a deep closed state of the pore

Remarkably, the R3Q at 0 Ca^2+^ resulted in a channel structure similar to WT apo, but with notable differences. We found a considerable narrower point in the inner cavities of R3Q apo (radius ∼4.2 Å, HOLE) when compared to the WT apo structure (radius ∼5.7 Å, HOLE) (**Figure 3B**). The hydrophobic environment of the permeation pathway was higher in the closed state, likely a consequence of sidechain rotations for residues I301, F304, and I308 toward the internal cavity (**Fig S10**). In particular, the side chains of I308 is the narrow point in the permeation pathway (radius ∼4.2 Å, HOLE) (**Figure 3 C-D**). Although I308 constricted the pore in R202Q apo, this closure may still allow the permeation of the hydrated K+ (radii 3.5-4.0 Å). However, a hydrophobic pore with a diameter <10 Å can undergo de-wetting, posing a large free energy barrier to ion conduction^38,39^. This structure represents the narrowest internal vestibule of a BK channel. We suggest that R3Q is able to stabilize a deep closed state of the BK pore and that conformation is evidence of an activation-deactivation gating mechanism based on the reorientation of the S6-based inner bundle helix. This mechanism appears to be conserved in other RCK-containing Ca^2+^ dependent K+ channels as recently suggested for MthK^40^.

In the BK channel, quaternary ammonium (QA) blockers may access the central cavity in the open and closed state of the channel^28^, unlike other Kv channels^41^. but, bbTBA, a bulky derivative of tetrabutylammonium, does not block the channel in a complete state-independent fashion^30^. Functional evidence suggests that bbTBA blocks the channel in an intermediate close state, but the fully closed BK channels are not sensitive to block by bbTBA^30^. We tested whether the geometry of the pore allows the entry of bbTBA to the internal cavity. We found that the permeation pathway observed in other BK structures^5,6,10,25–27^ at 0 Ca^2+^ conditions are large enough to allow bbTBA access to the blocking position, but the tight closure at I308 was sufficient to occlude the entry of bbTBA in R3Q apo (**Fig S11**). This suggests that previous BK apo structures likely correspond to a partially close state. However, when the VSD is firmly in its resting state (R3Q apo) the channel finally adopts its fully closed conformations. We submit that this structure is conformationally equivalent to that of the close BK channel at negative potentials.

Previous MD simulations ^42–44^ support the idea that the barrier for ion permeation in the closed state results from the de-wetting that occurs within the constraints of a narrow hydrophobic pathway. Similar mechanisms have been proposed in other channels^45–47^. We performed MD simulations of the pore domain using the R3Q closed-state and WT apo structures. We restrained the movement of the α-carbons of the S6 helix to determine whether R3Q is in a closer state of the pore than the WT apo structures. We found that in two different classes of R3Q apo structures, fewer water molecules are stabilized in the internal cavity (**figure S12**). When the structural constraint is removed, the cavity becomes devoid of water and lipids start to populate the inner cavity using lateral windows.

### Membrane-facing fenestrations in the closed channel

Rotation of the S6 helix required to close the channel not only exposes its hydrophobic surface to water in the conduction pathway (**figure S13**) but also opens membrane-facing fenestrations in the closed structures. These fenestrations also appears in other BK structures in apo condition, but not in Ca^+2^-bound structures^5,6,10,25,26^. They have been identify as alternative pathways for hydrophobic molecules to reach the inner cavity, in both MthK^40^ and dSlo1^26^. The membrane-exposed windows in the inner cavity allow for lipid binding and modulate the probability of channel opening and could explain existing data pointing to cholesterol stabilizing the closed state of BK. However, the ability of lipid molecules to populate the inner cavity might be an integral part of channel gating, and a condition of equilibrium reached during the purification of the protein ought to be tested experimentally. This phenomenon raises interesting questions regarding the process of rehydration of the inner cavity during channel opening.

### Resting VSD produces disown interaction between VSD and CTD

When compared to WT -apo, we observed small conformational changes in the CTD of the fully closed R3Q-apo. The most relevant difference was found in the αB helix, which is broken at position L376 (**Figure S14**). Whereas the αB helix of WT – apo span from L372-R382, similar to all our open structures. An artificially broken αB helix produced by the introduction of proline in the position hL380 (aL376), decreases both voltage and Ca^2+^ activation^48^. The resemblance in the αB helix structure of hL380P and R3Q apo suggests that the complete span of αB helix (L372-R382) is essential for the efficient coupling of CTD-VSD.

According to the available data, the deep closed state stabilized by mutant R3Q in the apo condition, appears to be equivalent to the fully closed state of BK in the presence of a resting potential. The introduction of Q202 likely forces the voltage sensor into its state, as it cannot occupy the hydrophobic center in a stable condition. However, despite this, the R3Q apo structure still manages to transition to the open state of their channel, suggesting that some form of coupling mechanism remains intact in this structure.

### Conformational Changes in the Pore During the Closed to Open Transition

We found open channel structures in VSD active-apo and VSD resting Ca^2+^-bound conditions. These open structures have similar structural elements. At the selectivity filter, four K^+^ ions were found to occupy the coordination sites (S1-S4). The internal cavity displayed a clear density in the middle of the cavity, surrounded by eight densities, potentially representing a potassium ion in the center of the cavity surrounded by water (**figure S15**). S6 adopts an open conformation in the R1Q (apo-and Ca^2+^-Bound), R2Q (apo-and Ca^2+-^Bound), and R3Q Ca^2+-^bound structures (**Figure 3A**). The pore-lining helix S6 tilted at position I299, moves upward (6.7 Å at Cα 315), resembling the position of the Ca^+2^-bound WT structure (5TJ6). We identified a stable electrostatic interaction between E310 (S6) and R223 (S5) in open structures. In addition, three putative π-interactions were identified. In the upper part of S6, the aromatic rings of F296-F300 face each other almost in a parallel conformation with a distance of ∼ 4 Å (**figure S16**). In the inner cavity, F304 form a likely π-cation with carbonyl nitrogen from S275 backbone (∼4.3 Å), whereas F307 formed a parallel stack with F231 (∼4 Å) in S5. The importance of hydrophobic interactions in BK S6 has been previously reported, particularly the alanine substitution of hF315 (aF304), which produces stable closed channels under Ca^2+^-free conditions^23^. In addition, MD simulations have found that aF304A plays an important role in closing the channel^42^. We identified two distinct open-channel structures, each activated by separate stimuli, each achieving an open state independent of calcium: the R1Q-apo structure, which is voltage-activated, and the R3Q-Ca^2+^-bound structure. This finding allows us to evaluate the structural features necessary to reach the open state and to investigate the potential coupling mechanisms facilitating channel opening.

### Calcium independent opening

The mutant R3Q led to an open channel under Ca^2+^-bound conditions, even with the voltage sensor in its resting state. These CTDs are nearly identical to previous structures in *Aplysia* and *human* BK (**figure S17**) ^5,6,8,9^. In the Ca^2+^ bowl, located in intersubunit interface, the Ca^2+^ ion is coordinated by residues D905, D907, G899, and D902 on the RCK2 domain of one subunit. Whereas, Ca^2+^ binding site located at RCK1 is coordinated by D356, E525, R503, G523 and E591 (**figure S7**) in all our Ca^2+^-bound structures. All these residues have been previously identified in mutagenesis studies as critical for Ca^2+^ binding at this site^49–53^.

Compared to the R3Q-apo structure, which shows few interactions between the CTD and VSD, the R3Q-Ca^2+^-bound structure reveals a robust set of interactions between these domains. Specifically, the carboxyl side chain of E377 in the αB helix of the CTD forms a hydrogen bond with the hydroxyl side chain of S219 (2.7 Å) in the S4-S5 loop of the VSD (**Figure 4B**). Furthermore, positive residues within the αB helix, including K381 and R382, form hydrogen bonds with D208 and T211 located at the end of S4. Additionally, residues at the Mg^2+^ binding site, such as N161 from the VSD, come into close contact with E363 and E388 at the CTD, coordinating Mg^2+^.

**Figure 4-.**
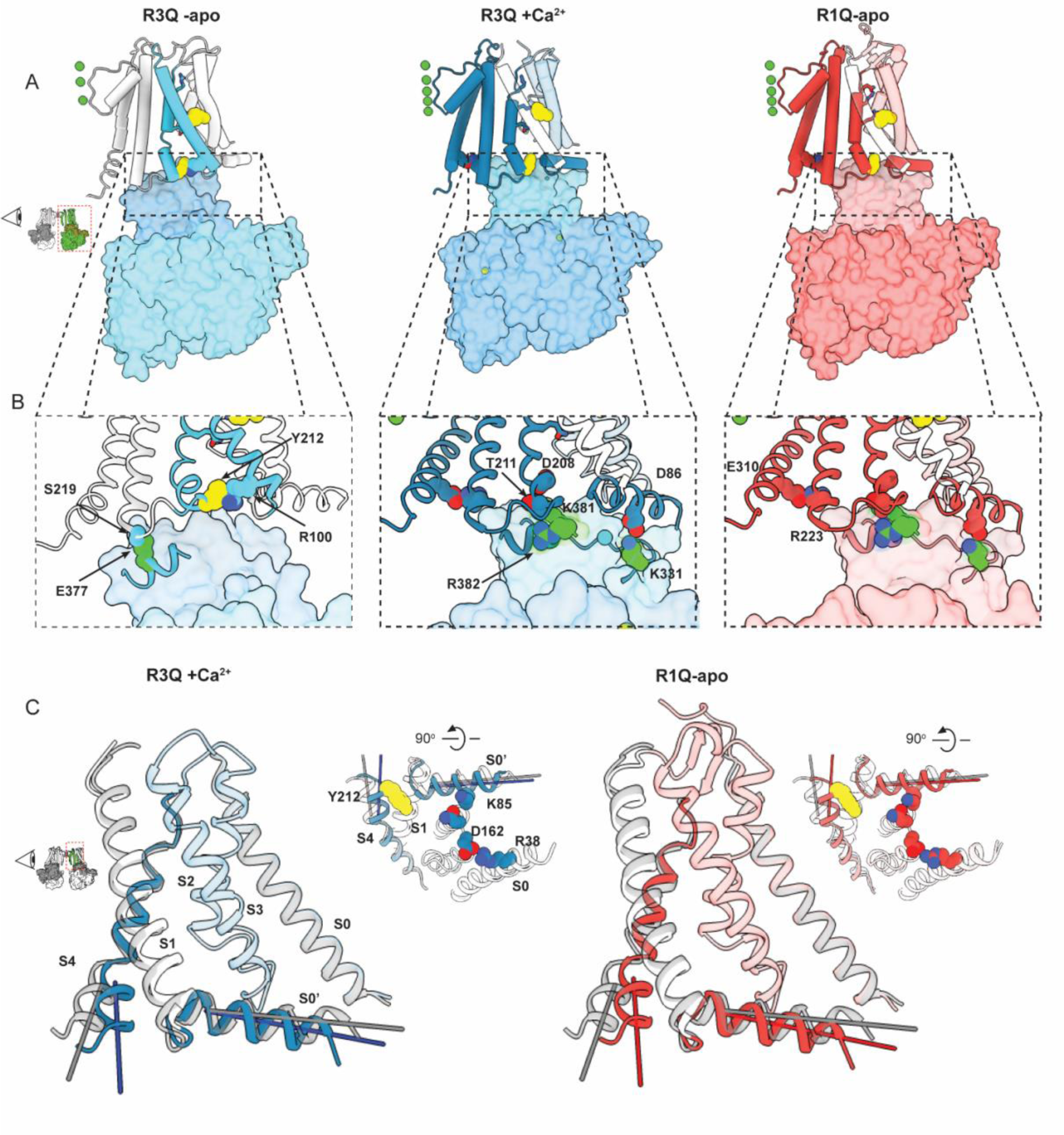
Reciprocal relation between VSD-CTD. (A) Side view of the VSD and their closest CTD. VSD is represented in a cartoon, whereas CTD is represented as a surface. Segments that are structurally coupled where colored and non-interacting segments are in white color. (B) Amplification of the interaction surface, where the segment of CTD that forms interaction with VSD is represented as a carton. For R3Q-apo the interacted segments were (C) Side view of the VSD of R1Q -apo and R3Q Ca^2+^-bound. Segments S0’-S4 are shown in cartoon representation. Segments S0’ to S3 are shown in transparency. The resting state VSD is shown in grey.

In the R3Q-Ca^2+^-bound structure, we observed that K331 in the C-linker forms ionic interactions with D86 (3.2 Å) at S0’. These interactions result in conformational changes at the bottom part of the S4 helix, which rotates from the M204 α-C by about 10°. Meanwhile, the S0’ rotates about 8° toward the cytoplasmic side, causing a change in the packing of the aromatic ring Y212. In the apo condition, the residue Y212 (S4) is packed between the backbone nitrogen of I209 and R100 (S1) between S4 and S1 (**Figure 4B-C**). However, in the Ca^2+^-bound state, Y212 is forming cation-π wit nitrogen backbone in S4 (I208) and S0’ (G94). The importance of Y212 has been highlighted by the alanine substitution at hF223A (aY212), which alters voltage sensitivity by increasing the channel open probability at negative voltages and decreasing it at positive voltages. Notably, the channel does not reach maximal activation without calcium, indicating that the interaction between S4 and S0’ may be critical for channel opening^24^.

### Voltage independent opening

Our VSD active structures (R1Q and R2Q) show that residues that participate in the coordination of Mg^+2^ are involved in the docking of the CTD VSD in the apo condition. Mutational analysis have identify residues on the interface between the VSD (D99 and N172 in BK) and the CTD (E374 and E399 in human Slo1) important in the coordination of Mg^2+ 52,54,55^. early structures shown that this binding pocket is conserved in aplysia, where residues E363 and E388 (E374 and E399 in human) in the CTD, as well as N161 (N172 human) in the VSD coordinate the ion binding^5,6^. Moreover, the main functional effect of Mg^2+^ is the coupling of the VSD^55–57^.

in the R1Q apo, the displacement of the gating charge R2 couples the S4 with S2, bringing the segment S0 closer to the extracellular side of the VSD. This results in changes at the cytoplasmic end, where the S0 helix’s R38 forms hydrogen bonds with D162 (2.8 A) at the S2-S3 loop, and N161 contacts the S0’ segment by forming a hydrogen bond between N161 and K85 (2.8 A). This causes the S0’ segment to rotate about 8° toward the cytoplasmic side, similar change found in the R3Q-Ca^2+^-bound structure (**figure 4B**). This conformation allows for the docking of the CTD. We found that K331 in the C-linker forms ionic interactions with D86 (3.2 Å) at S0’ is the main interaction that coordinating the coupling between sensors CTD-VSD. Consistent with mutagenic studies that show that alanine substitution at this position (hD99A) prevents the Mg^2+^ binding^55^. Analysis of the binding using the allosteric model predict that Mg^2+^ binding is state depend, it increases P_O_ only when voltage sensors are activated ^56^. The distance between residues N161 and E388 observed in the R1Q (4.3 Å) and R2Q (4.7 Å) variants under apo conditions, is like the distance the have WT Mg^2+^ (4.3 Å between N161-E388) bound structure, but without the ion in its binding site the electrostatic interaction between these residues seems unlikely. The final result of the formation of hydrogen bonds between S0’ D86 and K331 at the C-linker (3.2 A), is the conformational change between S0’ and S4 via Y212 (**figure 4B**). These findings indicate that the primary pathway for channel opening resides in the protein’s intracellular interface, and that electrical and chemical stimuli activate both sensors reciprocally, utilizing a common interface to couple the sensors and leveraging similar energy pathways to open the channel.

### A Non-canonical pathway connect voltage sensor with the pore

The propagation of energy transduced by the BK sensors to the pores follows a common pathway. All our open structures show that BK sensors contact the pore via steric interaction between the αB helix of the CTD and loop S4-S5, but also the C-linker and segment S0’(**figure 4B-C**). Their contact results in the formation of hydrogen bonds between R223 and E310, which appear to be canonical interaction-coupling sensors and pores identified in our open structures. Mutations in either R223 or E310 reduce coupling energy to such an extent that coupling cannot be measured without calcium24. This electrostatic interaction S5-S6 appears in all open channel structures (**Figure 4**). However, alanine introduction at the homologous position 310 (E320A) also decreased basal activity, indicating that it also stabilized the closed state. at the open pore of BK, segment S6 adopts an alpha-helix configuration compared to our closed structure, where the deep S6 helix (from position 308 to 318) gains flexibility around P309, which may allow the rotation of hydrophobic residues toward the conduction pathway (**figure 3**). This suggests that pore opening is achieved by straightening the passive spring that connects the TM and the cytosolic domain^58^. calcium and voltage can influence this spring through steric contact, which increases the stability of segment S6 via steric interactions^24^. However, these mechanical models do not fully explain the channel gating.

We observed state-dependent packing of hydrophobic residues in open structures (**figure S16**). This correlates with the decrease in hydrophobicity in the open state as well as with the closing of the membrane-facing fenestration (**figure S13**). However, in the closed state, these residues did not interact with each other. Evidence from MthK suggests that lipids may occupy the inner cavity in the closed state using fenestrations as the permeating pathway^40^. This suggests that the channel may use the energy from the sensor to close the fenestration, thereby restoring the hydrophilic environment permeation pathway. The introduction of charged residues in S6 segments results in permanently open channels or a leftward shift in the G-V relationship^59,60^. To test the idea that hydrophobic interactions stabilize the open pores, and that their stability is a function of the energy coupling, we solved the structure of F304A in activated Ca^2+^-bound conditions. Alanine replacement at this position in hBK (hF380) produces channels that open with low Po even at high Ca^2+^ concentration^23^.

We found that the structure of the F304A Ca^2+^-bound channel is in an open state, similar to the R3Q Ca^2+^-bound channel. The voltage sensor of F304A was resting in the Ca^2+^-bound condition (**Figure 5A**) without the electrostatic constraint imposed by the R3Q mutation. Importantly, we found that the positions of the charged residues resembled those in the R3Q mutation, corroborating the small transition of the VSD charged residues. The residues interacting in the cytoplasmic end of the TM region in open channel R1Q apo and R3Q Ca2+-bound are also found in F304A with Ca^2+^ binding (**Figure 5A**). Interestingly, the fenestrations observed in R1Q-apo were not fully closed in activated F304A channels (**figure 5B**). Surprisingly, the structure of the S6 segment was also affected by alanine replacement at 304 (**figure 5B**), but the hydrophobic residues F304, I308, and A312 that close the fenestration in structures, the open channels R1Q-apo, and R3Q-Ca^2+^-bound (**figure 5C**) were in the same position, except for A304, which has a side chain pointing to the internal cavity. This suggests that the internal cavity of open F304A has a small, but open fenestration in its Ca2+-activated conformation. hydrophobic interactions found in open R1Q-apo underwent positional changes of the side chain in the R3Q Ca2+-bound channels, which are well defined in the density map (**figure 6A**). Since R3Q Ca2+-bound channels must have a lower open probability than R1Q-apo at 0 mV, we expected to find a correlation with activated F304A channels. Indeed, the Ca2+-bound structure of the hydrophobic residues coupling S5-S6 in R1Q-apo does not align with either F304 or R3Q(**figure 6B**). We found that F304 likely formed a π-cation with carbonyl nitrogen from the S275 backbone (∼4.3 Å) in all open-channel structures. This interaction may be relevant for channel activation, since the mutation of tryptophan at this position facilitates channel opening. Surprisingly, both hF380A and hF380W mutations decreased the channel conductance by approximately 60% and 50%, respectively. This indicates that a factor other than the volume of the residue may be important in controlling channel conductance.

**Figure 5.**
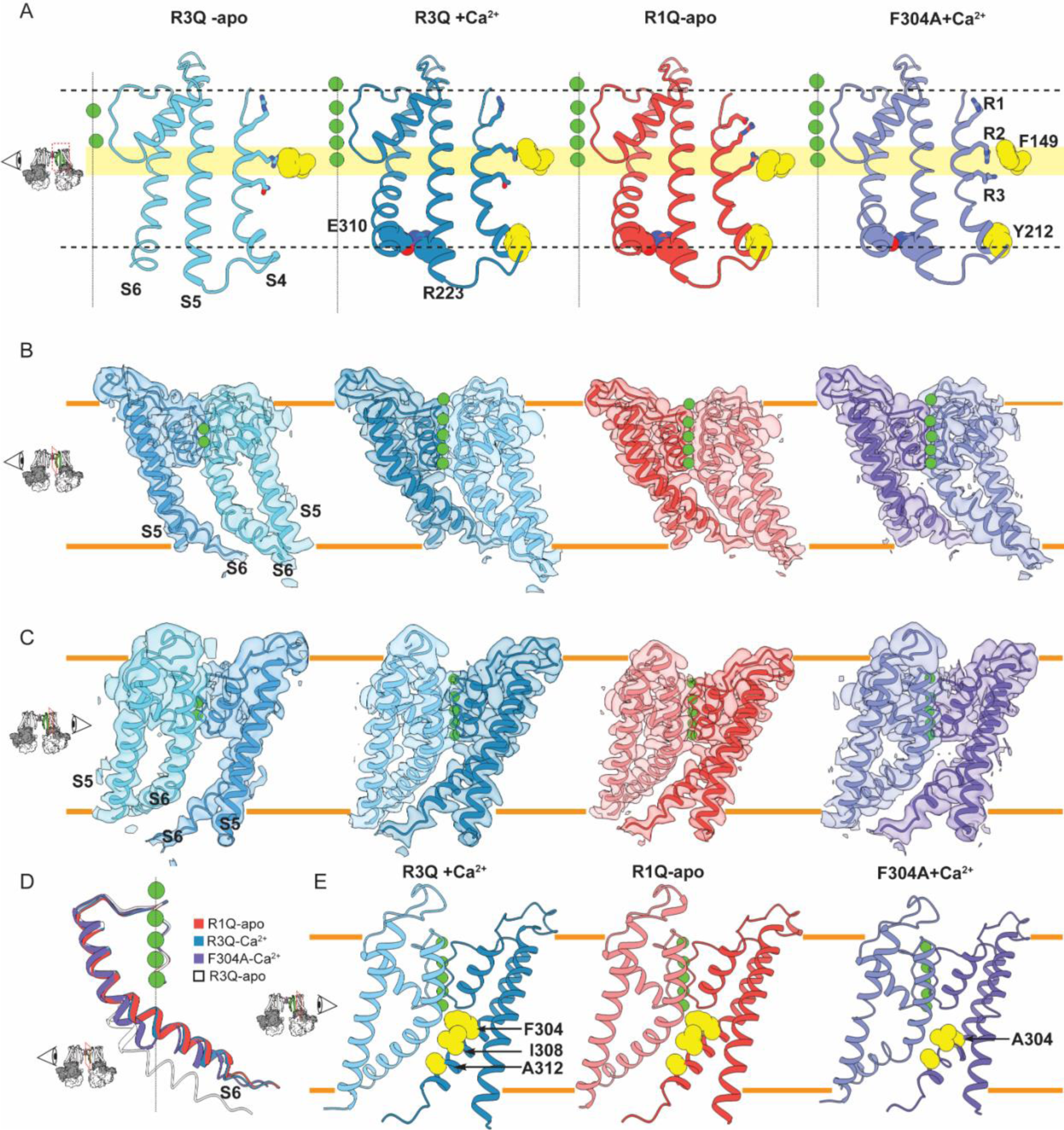
State-dependent membrane-facing fenestration. **(A)** Side view of VSD and PGD including segments S4-S6. The position of charged residues at S4(196,199, and 202) are in stick representation whereas the hydrophobic gasket (F149) and residues coupling segments S4(Y212), S5(R223), and S6(E310) are represented in vDW. **(B)** Internal Side view of PDG. Two contiguous subunit are shown in cartoon representation superimposed on their Cryo-em density maps. **(C)** Peripheral side view of PGD, including segments S5 and S6 superimposed with their Cryo-em density maps. **(D)** Internal view of the pore segment S6. **(E)** Peripheral side view of two contiguous PDG in cartoon representation. Hydrophobic residues F304, I308, and A312 are shown in vDW representation.

**Figure 6.-.**
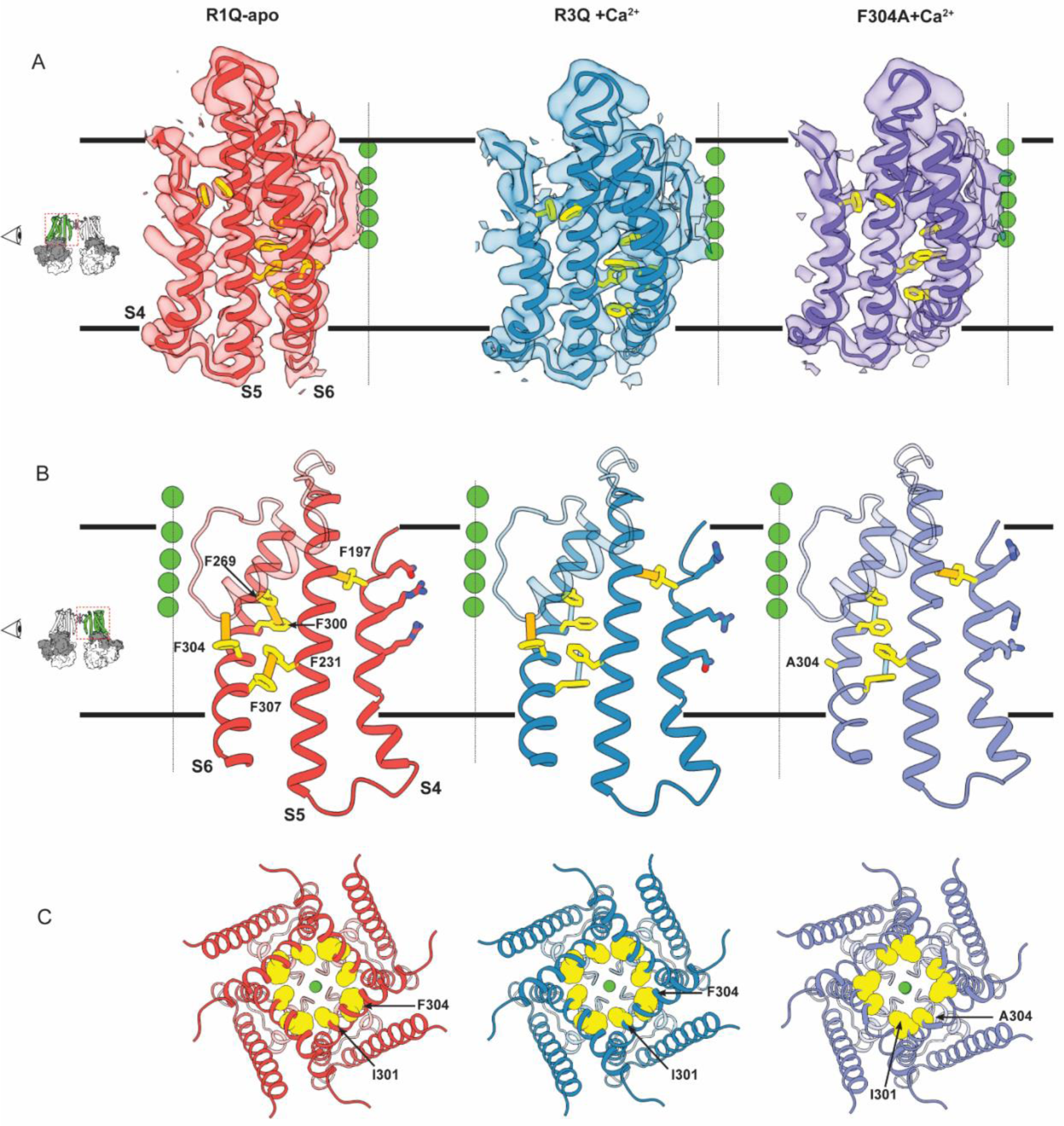
non-canonical coupling in TM domain. **(A)** Side view of TM segments S4 to S6, superimposed with their Cryo-em density maps. The residues forming hydrophobic interactions are shown in stick representation and color in yellow. **(B)** Hydrophobic interactions highlight by links connecting the center of aromatic ring of pair F296-F300 (S6), F231-F300 (S5-S6). Also pi-cation interactions between F304 and nitrogen backbone 275, F197 with backbone of F245 are shown in orange. Normal vector that are not forming hydrophobic interactions are shown in light blue. **(C)** Bottom view of the pore internal cavity showing the hydrophobic ring in the open state. Residues 304 and 301 are shown in vDW representation.

We found that F304 and I301 (hL377) form a hydrophobic ring in the open state (figure 6C). In the closed state, the side chains point toward the internal cavity. Our results agree with the structure proposed by MD simulation^23^, which predicts the formation of this ring in the open state. Early studies have predicted that F304 and I301 also stabilize the closed state^61^; however, recent MD simulations have predicted the role of these residues in pore de-wetting^42^. Our results agree with hydrophobic ring formation in the open state, but do not discard the role of its residues in the closed state. We found that conformational changes around F197 could be mechanically coupled with VSD movement to the pores (**figure 6**). The aromatic ring of F197 is packed between the backbone nitrogen atoms of S4 and S5 in both open and closed structures (**figure S18**). However, the distance between S4-S5 changes the open R1Q-apo compared to the low Po open R3Q and F304, which changed together with the reorientation of the side chain of several aromatic residues in both S5 (F241,F231) and S6 (F296,F300,304, and F307). The observed differences provide a structural depiction of possible non-canonical interactions by which the sensor and pore could be energetically linked. Small differences in Q-V were found in the hF380 mutant, suggesting that the energetic link between the sensor and pore was low. The conformational changes observed here and previous functional evidence^62^ agree that saturating Ca^2+^, the voltage sensor movement, increases Po mainly through non-canonical interaction, which increases the channel opening rates, with limited effects on the closing rates.

The propagation of the energy transduced for sensor of BK to pore follow a common pathway. All our open structures show that BK sensors contact the pore via steric interaction between αB helix of the CTD and loop S4-S5, but also the C-linker and the segment S0’(**figure 4B-C**). their contact result in the formation of hydrogen bonds between R223 and E310, that appears to be the canonical interaction coupling sensors and pore identify in our open structures. Mutations in either R223 and E310 reduce the coupling energy at such extent that the coupling cannot be measured in absence of calcium^24^. This electrostatic interaction S5-S6 appear in all open channel structures (**Figure 4**). But alanine introduction at homologous position 310 (E320A) also decreases the basal activity indicating that also stabilize the closed state. at open pore of BK, segment S6 adopts a alpha-helix configuration compared with our closed structure where the deep S6 helix (from position 308 to 318) gain flexibility around P309, which may allows the rotation of hydrophobic residues toward the conduction pathway (**figure 3**). Which suggest that the opening of the pore is achieved by a straighten of the passive spring that connect TM and cytosolic domain^58^. Ca^2+^ and voltage can influence this spring by steric contacts that increase the stability of the segment S6 via steric interactions^24^. However, these mechanical models not fully explain the channel gating.

We observed state dependent packing of hydrophobic residues in open structures (**figure S16**). This correlates with the decrease in hydrophobicity in the open state, but also with the closing of the membrane-facing fenestration (**figure S**). whereas in the closed state these residues are not interacting. Evidence from MthK suggest that lipids may occupy the inner cavity in the close state, using fenestrations as permeating pathway^40^. This suggest that the channel may using the energy from the sensor to close fenestration restoring hydrophilic environment permeation pathway. Introduction of charged residues in S6 segments result in permanently open channels or leftward shift in the G-V relationship. To test the idea that hydrophobic interaction stabilizes the open pore, and their stability is a function of the energy coupling we solve the structure of F304A in activated Ca^2+^-bound condition. Since the alanine replacement at this position in hBK (hF380) produces channel that open with low Po even at high Ca^2+^ concentration^23^.

We found the structure of F304A Ca^2+^-bound condition is open state like R3Q Ca^2+^-bound channel. The voltage sensor of F304A was in resting in Ca^2+^-bound condition (**Figure 5A**), without the electrostatic constrain imposed by R3Q mutation. Importantly we found the position of the charged resemble those in R3Q mutation corroboration the small transition of the VSD charged residues. The residues interacting in the cytoplasmatic end of the TM region in open channel R1Q apo and R3Q Ca^2+^-bound, are also found in F304A with Ca^2+^ bound (**Figure 5A**). Interesting, the fenestration observed in R1Q-apo are not fully closed in the activated F304A (**figure 5B**). surprisingly the structure of the S6 segment is also affected by alanine replacement at 304 (**figure 5B**), but the hydrophobic residues F304, I308, and A312 that close the fenestration in structures the open channels R1Q-apo, R3Q-Ca^2+^-bound (**figure 5C**) are in the same position except for A304, that has a side chain pointing to the internal cavity. This suggest that internal cavity of the open F304A has a small but open fenestration in its Ca^2+^-activated conformation. hydrophobic interactions found in open R1Q-apo experienced positional changes of the side chain in R3Q Ca^2+^-bound channels, that are well defined in the density map (**figure 6A**). since R3Q Ca^2+^-bound channels must have lower open probability than R1Q-apo at 0 mV, we expected to find a correlation with activated F304A channels. Indeed, the Ca^2+^-bound structure the hydrophobic residues coupling S5-S6 in R1Q-apo, does not align neither in F304 nor R3Q(**figure 6B**). we found that F304 form a likely π-cation with carbonyl nitrogen from S275 backbone (∼4.3 Å) in all open channel strcuttures. This interection may be relevant for channel activation since the mutantion for tryptophan at this position facililtes the channel opening. Suprunsingly, both mutations hF380A and hF380W mutations decrease the channel conductance about 60% and 50%, respectively. Indicating that a factor other that the volume of the of the residue may be important in controlling the channel conductance.

We found that F304 and I301 (hL377) form a hydrophobic ring in the open state (**figure 6C**). Whereas in the closed state their side chain are pointing to the internal caity. Our result agree with the proposed structure by MD simulation^15^ that predict the formation of this ring in the oen state. Early studies have predict that F304 and I301 stabilize the also the closed state^61^, but also recent MD simulation predict a role of this residues in the pore de-wetting^42^. Our result gree with hydrophobic ring formation in he open steate but not discard the role of tis residues in the close state. The non-canonical interaction connecting the pore an the VSD are between the pore and the voltage sensor it

**Figure 7.-.**
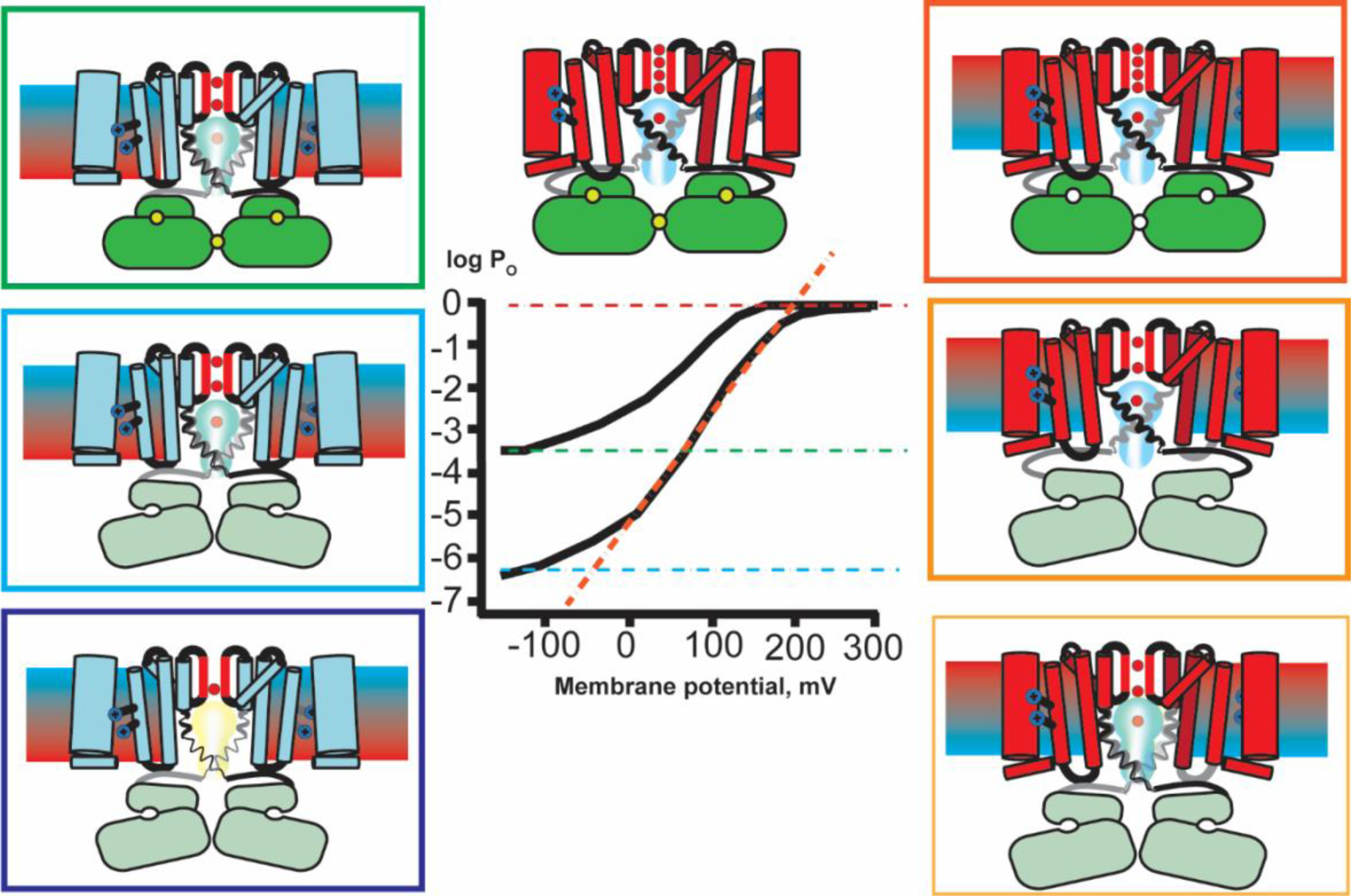
non-canonical coupling in TM domain. Schematic representation of two activation gating mechanisms. From the closed state at the bottom left corner, Ca^2+^ activation schemes from bottom to top. In the right side scheme represent the voltage activation.

## DISCUSSION

The role of the Ca^2+^ and voltage activated potassium channel (BK) as signal integrator in excitable cells has evolved as a consequence of the remarkable cooperativity between its voltage sensors and Ca^2+^-binding RCK domains. Although BK has long been characterized by its shallow voltage sensing and fast charge transfer kinetics, the molecular mechanism of voltage sensing and the nature of its coupling to the Ca^2+^-binding domains remain a matter of debate since its early characterization ^15,68^. Neutralizing the charged residues in the VSD has led to conflicting results ^31,32,71^. Whereas Ma et al., ^31^ suggested that the gating particles were distributed in S2, S3, and S4, Carrasquel-Ursulaez et. al ^32^ results indicated that the gating particles were contained in the S4 transmembrane segment R210 and R213). BK gating current behavior has also led to two contrasting interpretations regarding the degree of coupling with Ca^2+^ sensors: either little or no coupling ^15, 24^, or a Ca^2+^-driven modulation of the BK channel voltage dependence ^22,23^. BK structures in the apo and Ca^2+^-bound structures ^5,25^ also point to a strong coupling between VSD and Ca^2+^ sensors. Here, we show how the charged residues in S4 segment regulate the operational cycle of the voltage sensor. Neutralizing these residues stabilizes the active and resting states of voltage sensors. We successfully determined five high-resolution structures representing the open state, illustrating the independent mechanisms of voltage and calcium in the channel activation. In addition, we resolved two channel structures in the closed state. These structures reveal the rotation of the S6 side chains that form a hydrophobic inner cavity, open membrane-facing fenestrations, and narrow the conduction pathway. These structures, together with functional and computational analyses, contribute to a mechanistic understanding of the BK voltage dependence. They provide a plausible explanation for how the energy in the transmembrane electric field is transduced into pore opening, and furthermore, solve the lingering debate on the nature of the closed state of the BK channel.

### The operational cycle of the voltage sensor

We found that the VSD transition from the resting state to the active state involved a rotation of the side chain rotamer according to the expected direction of the electric field. Comparing the resting and active structures, no major vertical displacement was observed in S4. Additionally, the side chain of R196 exhibited minimal changes between the resting and active conformations of the VSD, confirming that the first arginine residue does not participate as a gating charge. However, R2 and R3 undergoes rotamer reorientation with a displacement of about 5 Å in their charged moieties (**figure 2A**). The charge movement is center around the hydrophobic gasket at the VSD. In the resting state structure, the electrostatic interaction between R2 and D175 maintains the connection between the S4 and S3 segments, whereas in the active state, R2 becomes exposed to the extracellular environment and interacts with D142, coupling the S2 segment. Meanwhile, R3, initially positioned in the intracellular space in the resting state, moved upward to occupy the hydrophobic center. This movement is effectively an electrostatic equivalent of the ‘one-click’ mechanism observed in phosphatases ^63,64^ and KAT1 channels^65^, channels in which the voltage sensor is coupled with a cytoplasmic structural element. But it takes place with minimal displacement of the S4 helix in the Z axis (perpendicular to the plane of the bilayer). This suggests a distinct mechanism of voltage dependence, where the S4 helix does not behave as a translating permion but net charge is translocated due to the rotamer reorientation of the active gating charges.

Our findings indicate that the majority of the distributed charged residues in S3 and S2 exhibit state-independent interactions, which serve to hold together the various segments of the VSD and facilitate the structural rearrangements that transmit VSD energy to the pore. This is particularly true for residues D175, R156, and E169 (hD180, hR167, and E186), which form distinct electrostatic interactions in both the open and closed conformations (figure S4). Therefore, the neutralization of these structural elements would result in a lack of coupling, as has been showed previously^31,32^. Our result support the idea that R2 (hR210) is the main gating charge. These residues displace outward from the hydrophobic center, where the electric field drops, and couple with segment S2, resulting in conformational changes at the cytoplasmic end of the VSD connected to the pore.

The degree of conformational change exhibited by the VSD of BK channels restricts their functional comparison with other ion channels. While Kv channels are capable of displacing a greater amount of charge (13 e)^66,67^, their kinetic movement of the voltage-sensing domain (VSD) is slower than the rapid transient movement of BK channels^68^. This difference in electromechanical mechanism between Kv and BK channels suggests that the gating mechanism in Kv channels differs from the mechanism found in BK channels.

### Coupled rearrangement among Voltage and Ca^2+^ sensor

We have found structural evidence that the activation energy transmitted by voltage and calcium utilizes the same conformational pathway to bias pore opening. It involves conformational changes in individual sensor molecules that affect each other through allosteric interactions. These interactions result in the energy in the transmembrane electric field being transduced through small sequential changes within the transmembrane segments, specifically S0’ to S4 to S5 to S6. Our findings align with previous functional characterizations, showing that the degree of voltage sensor activation modulates the apparent affinity for calcium^18^, and that the internal calcium concentration influences the activation energy and mid point of activation in the VSD^2^. Furthermore, even transient conformational changes in an individual sensor appear to be coupled. This is demonstrated through methods such as fluorometry, which tracks changes in the S4 region in response to transient pulses of calcium^21^, or the movement in the CTD following transient activation of the voltage sensor at negative voltage^19^.

We also report the opening of the channel by the exclusive activation of the calcium sensor, as predicted by allosteric models. However, we forced this scenario by locking the resting state of the VSD. We expected to find a configuration in which the VSDs are active, but the channel is closed for the mutant R2Q since this mutation impairs the coupling to the pore^32,36^. It is possible that the pore of *Aplysia* BK might have a facilitated pore because it has a different distribution of positive residues in the C-linker compared to hBK; in particular, the hK330 is changed by glycine. The alanine modification of this residues in hBK produces a 100-fold increase in the spontaneous opening of the pore^24^. Nevertheless, as our structures are in equilibrium, we cannot discard the possibility that the channel can be exclusively opened by conformational changes in the VSD with the calcium sensor in a contracted conformation. In summary, our findings provide evidence for a complex relationship between voltage and calcium sensors, highlighting a refined allosteric mechanism that is characterized by delicate conformational modifications in one domain leading to substantial functional shifts in the channel. This enhances our comprehension of sensor-pore interactions and offers a more precise framework for analyzing the dynamic regulation of ion channels.

### Conformational Change in the Pore

Minor alterations in the deep pore S6 helix, particularly the R3Q mutation, have a significant impact on the pore size. This is clearly demonstrated when comparing the R3Q closed structures with the wild type (WT) in its apo state. The R3Q mutation stabilizes the resting state VSD and concurrently induces a conformational change in the pore. This change leads to the exposure of hydrophobic residues (F304, I308, A313) along the S6 helix to the ion conduction pathway, thereby constricting the pore diameter. The narrowest constriction occurs at I308 (radius 4.2 A).

Molecular simulations have predicted a progressive de-wetting of the hydrophobic cavity when their diameter reaches less than 10 Angstroms^38^. This mechanism was previously postulated in human^43^ and Aplysia BK^42^ channels and our study provides the first structural evidence supporting this permeation mechanism in BK channels. This mechanism differs from Kv channels, where a cytoplasmic gate occludes the permeation pathway^41^. In BK channels, there is no evidence of a cytoplasmic gate as hydrophobic ions can block the channel in its closed state^28^. Notably, an artificial peptide resembling the inactivation ball of Shaker Kv acts as an open-channel blocker^69^, suggesting a change in the hydrophobic nature of the closed pore.

The geometric constraints imposed by the blocker bbTBA, with a diameter of 10 Å, lend credence to the notion that the R202Q-apo structure represents a more closed channel. Experimental evidence suggests that bbTBA can block channels in a semi-closed state, where at least one voltage sensor is active. However, fully closed BK channels are insensitive to bbTBA blockage^30^. Recently, another pathway for the entry of hydrophobic ions and drugs was suggested for MthK^40^, in which hydrophobic blocker reach the pore trough’s membrane-facing fenestration. These fenestrations are present in several BK channel structures, where lipid or detergent density appears to partition into.

The exact mechanism for opening the channel remains unclear. The presence of a double glycine motif in the S6 helix of hBK has been proposed as the point of flexibility when the helix bends. However, this motif is not conserved in other species, yet the channel still opens at the same point in structures from different species. Additionally, the occupation of the pore by hydrophobic ions and interaction with membrane lipids result in further conformational changes in the geometry of the pore. All of these observations suggest that the closed channel is highly heterogeneous and sensitive to physico-chemical properties of the surrounding membrane. This observation is reminiscent of the early definition of allosteric proteins that exist in two different conformational states, designated as ‘R’ (for relaxed) or ‘T’ (for tense) states^70^.

